# Hematopoiesis at single cell resolution spanning human development and maturation

**DOI:** 10.1101/2021.08.25.457678

**Authors:** Hojun Li, Jideofor Ezike, Anton Afanassiev, Laura Greenstreet, Stephen Zhang, Jennifer Whangbo, Vincent L. Butty, Enrico Moiso, Guinevere G. Connelly, Vivian Morris, Dahai Wang, George Q. Daley, Salil Garg, Stella T. Chou, Aviv Regev, Edroaldo Lummertz da Rocha, Geoffrey Schiebinger, R. Grant Rowe

## Abstract

Hematopoiesis is a process of constitutive regeneration whereby hematopoietic stem and progenitor cells (HSPCs) replenish mature blood cells. During maturation and aging, HSPCs shift their output to support the demands of prenatal development and postnatal maturation both at homeostasis and in response to stress. How HSPC ontogeny changes throughout life is unknown; studies to date have largely focused on specific individual ages, particularly at single cell resolution. Here, we performed single cell RNA-seq of human HSPCs from early prenatal development into mature adulthood. We observed shifts in HSPC transcriptional states and differentiation trajectories over time. We identified age-specific gene expression patterns throughout human maturation and developed methods for identifying, prospectively purifying, and functionally validating age-specific HSC states. Together, our findings define the temporal maturation of human HSPCs and uncover principles applicable to age-biased blood diseases.

**Summary:** Single cell RNA sequencing reveals that the mechanisms of human hematopoietic stem and progenitor cell (HSPC) fate commitment change over a lifetime from gestation to mature adulthood.

## Main Text

Over a lifetime, stem cells must continuously adjust their prioritization of self-renewal and differentiation to execute tightly choreographed programs of development and maturation (*1*). Although stem cell ontogenies have been finely mapped at specific stages of human life, how stem and progenitor cell ontogeny changes over the course of a lifetime remains a fundamental, but unanswered question (*2-9*). To meet age-specific physiologic demands, the human hematopoietic system adjusts the output of mature cells capable of oxygen transport, hemostasis, innate and adaptive immunity, and tissue repair; however, the molecular regulation of these age-dependent alterations is not well understood.

Since lineage destiny is transcriptionally encoded within HSPCs, we performed single cell RNA-seq (scRNA-seq) of human HSPCs to reconstruct hematopoiesis over the course of human maturation. We procured CD34^+^ HSPCs at 12 time points across human development, from 10 weeks post-gestation to 45 years of age (1-2 donors per time point) and used the InDrop platform to profile individual transcriptomes (**Fig. 1A**)(*10*). After filtering of doublets, non-hematopoietic cells, and cells with poor coverage (see methods), we retained 38,873 HSPCs with high quality profiles. We performed dimensionality reduction with principal component analysis (PCA), followed by embedding via Universal Manifold Approximation and Projection (UMAP) and partition-based graph abstraction force directed layout (FDL) to visualize individual ages and the overall dataset (**Fig. 1B,I, fig. S1**)(*11, 12*). Expression of lineage specific marker genes revealed clear transcriptional priming of lymphoid and myeloerythroid lineages (**Fig. 1B-H, fig. S2**)(*8, 13, 14*).

**Figure 1.**
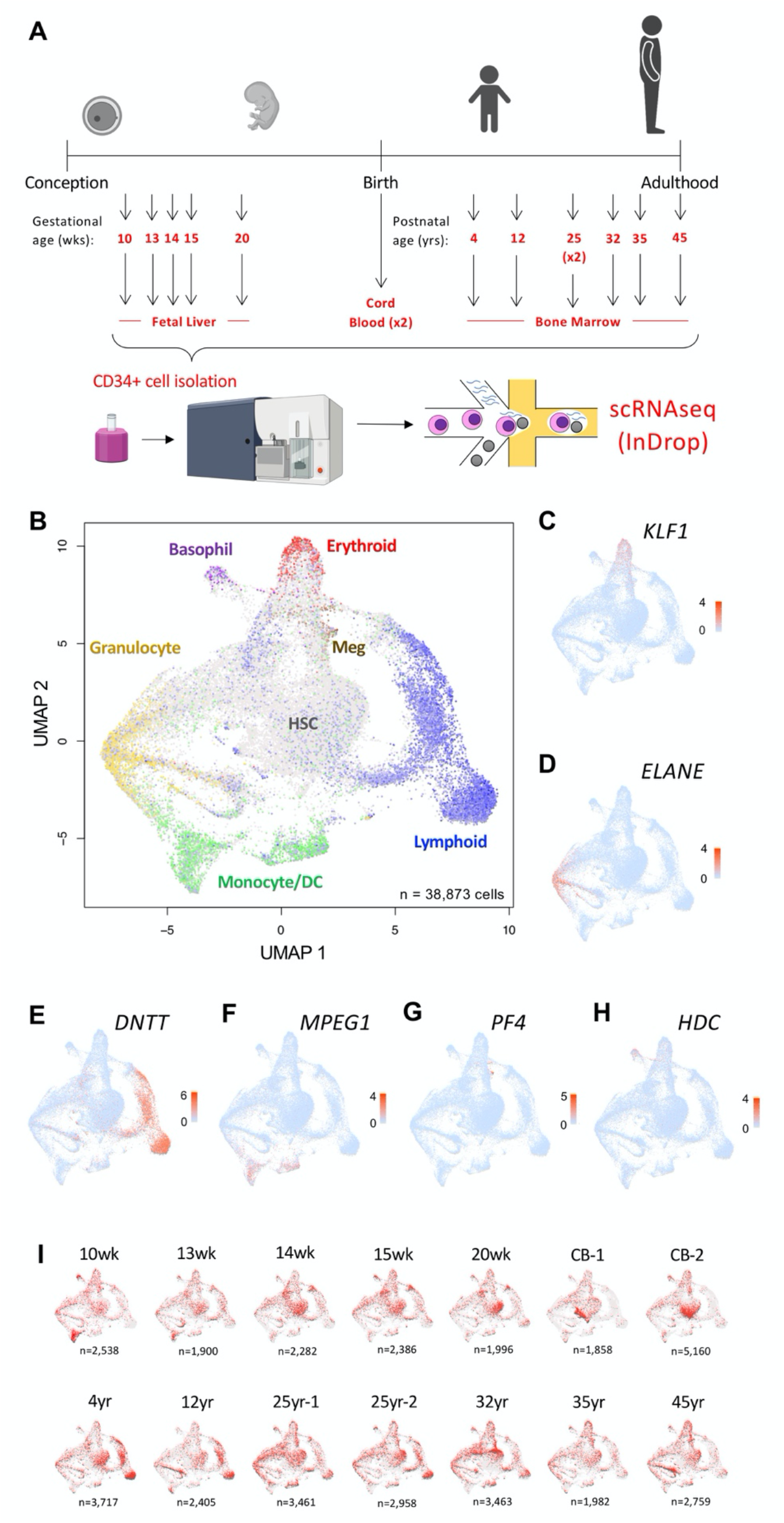
Single cell transcriptome profiling of human HSPCs from fetal life through adulthood. **(A)** Schematic of HSPC sampling. **(B)** UMAP-based embedding of all cells profiled, highlighted based on lineage scores for each individual cell. **(C-H)** Relative expression of lineage marker genes. **(I)** Sample distribution across the UMAP embedding for specific time points (red) relative to all other samples (gray).

We identified 21 distinct HSPC states by unsupervised clustering (see methods) that vary in abundance over time but are detectable at all time points (**Fig. 2A,C,F, fig. S3**). We used the singleCellNet algorithm to score each cluster against an existing human adult bone marrow scRNA-seq dataset (**Fig. 2B**)(*14, 15*); examining them in the context of the FDL and UMAP, we hierarchically ordered and annotated HSPC states spanning from the most undifferentiated HSCs and multipotent progenitors (MPPs) to unilineage-restricted progenitors (**Fig. 2A**)(*12*). Cells in HSC and MPP states were located centrally in both UMAP and FDL space with branches terminating in lineage-restricted progenitors (**Fig. 2A, fig. S3**).

**Figure 2.**
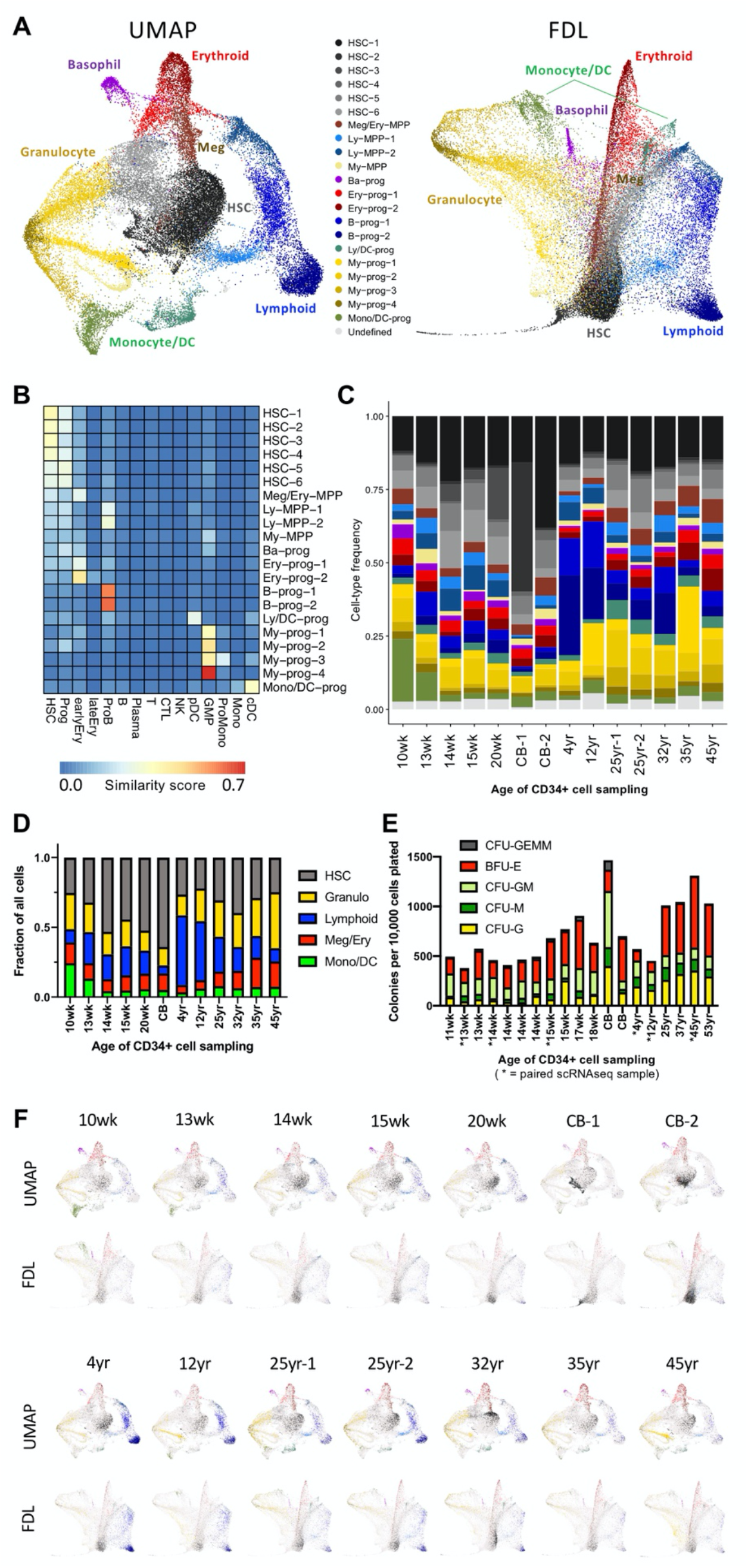
Unbiased clustering identifies lineage biases of human HSPCs. **(A)** HSPC subtypes from Louvain clustering displayed on UMAP and FDL. **(B)** Classification of HSPC clusters based on similarity to bone marrow cell types using SingleCellNet. **(C)** Barplot of distribution of individual HSPC clusters over human life. **(D)** Barplot of distribution of general progenitor cell types over human life. HSC clusters (HSC1-6), granulocyte clusters (My-MPP, Ba-prog, My-prog-1-4), lymphoid clusters (Ly-MPP-1-2, B-prog-1-2), Meg/Ery clusters (Meg/Ery-MPP, Ery-prog-1-2), mono/DC clusters (Ly/DC-prog, Mono/DC-prog). **(E)** Colony formation of CD34+ cells from human fetal liver and bone marrow. **(F)** Distribution of HSPC clusters over human life displayed on UMAP and FDL.

We found that the abundance of HSPC states and usage of differentiation trajectories changes over time. Early gestation fetal liver (FL, 10-13 weeks) prioritized trajectories toward myeloid output; later in gestation prioritization shifted toward HSC expansion (**Fig. 2F, fig. S4**). Erythroid production was maintained throughout gestation, supportive of growth within the hypoxic uterus (**Fig. 2C,D,F, fig. S4**). In childhood, lymphoid progenitors are relatively abundant, but diminish with age; myeloid and erythroid output increases over time as recapitulated in clonogenesis assays (**Fig. 2C-F, fig. S4**). The pediatric wave of lymphopoiesis occurs via differentiation through the Ly-MPP-2 state, which diminishes with age, while Ly-MPP-1 maintain steady-state lymphoid output throughout life (**Fig. 2C,F, fig. S4**). These overall patterns of lineage prioritization are supported by changes that occur in peripheral blood leukocyte populations across human life and the myeloid-biased hematopoiesis of mature adulthood (*16, 17*).

We next investigated the processes of age-specific lineage commitment using both Population Balance Analysis and an adaptation of Optimal Transport to systems in equilibrium (*18-20*). Both approaches model cell trajectories with the same diffusion-drift equation, which represents Waddington’s landscape, and both produce the same output: an estimate of the lineage commitment of each cell, for each lineage (i.e. lineage fate probabilities). While these two methods differ in their underlying mathematical approaches, they produced reassuringly similar results. In both approaches we found high myeloid differentiation probability for early gestation fetal liver HSPCs, with a mid-gestation shift toward more balanced hematopoiesis; higher lymphoid probability predominated in childhood, shifting back to higher myeloid probability in adulthood (**Fig. 3A, fig. S5**). These analyses also distinguished age-biased fate probabilities within the HSC state subsets that appeared uncommitted in UMAP and FDL space (**fig. S6**).

**Figure 3.**
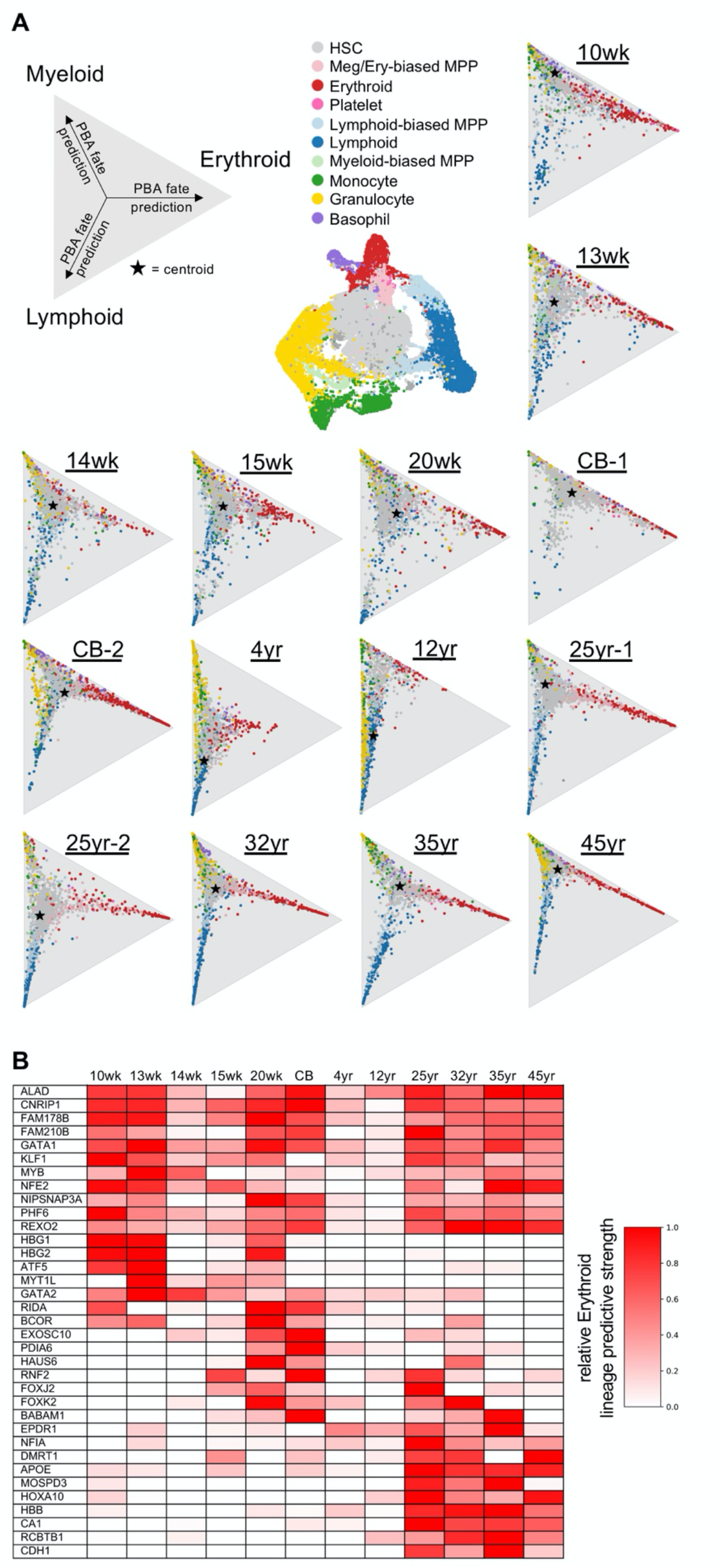
Single cell lineage fate and lineage gene marking from fetal life through adulthood. **(A)** Individual cell fate probability of erythroid, myeloid, and lymphoid differentiation based on population balance analysis (PBA). Proximity of each cell dot to each vertex indicates likelihood of differentiation toward that lineage. Centroid indicates average fate of all cells in that specimen. **(B)** Relative strength of select individual genes predicting erythroid commitment at different timepoints.

To identify the mechanisms underlying changes in lineage commitment during human development and maturation, we searched for transcripts whose levels are associated with age-specific differentiation. Elastic Net semi-sparse regression of cells undergoing lineage commitment identified transcripts for each lineage whose levels either consistently predicted fate probabilities, or only predicted fate probabilities at certain ages (*21*). For each lineage, we identified the top 200 transcripts showing either consistent or age-specific prediction of fate commitment (**Table S1**). For example, during erythroid differentiation we observed expected expression patterns in temporally consistent (*GATA1, KLF1*) and variable (*HBG1, HBG2, HBB*) erythropoietic genes, as well as several novel age-associated factors (**Fig. 3B**). These data indicate that while certain factors control differentiation throughout life, differentiation programs are also modified with age.

In addition to age-dependent programming of normal hematopoietic development, many types of leukemia are biased toward certain ages (*14, 22, 23*). We used singleCellNet to classify blasts of childhood B-acute lymphoblastic leukemia (B-ALL) and adult acute myeloid leukemia (AML) profiled by scRNA-seq (**fig. S7**)(*14, 22*). In childhood B-ALL, we observed shifting classification from diagnosis to relapse, with several cases adopting more differentiated states, and notably one case recruiting the fetal-biased HSC-3 state (**fig. S7**). In poorly differentiated adult AMLs, activation of differentiation programs occurred with treatment (**fig. S7**). These data demonstrate potential application of our atlas of human HSPC maturation to pinpoint leukemic differentiation state, uncover activation of oncofetal programs, and novel potential mechanisms of therapeutic evasion via differentiation or dedifferentiation.

As age-dependent changes in stem cell states have yet to be described at single cell resolution in humans, we sought to determine whether such a phenomenon occurs in HSCs by querying for age-specific HSC states that have yet to be detected in prior seminal studies of narrow developmental time windows (*24-26*). We observed that the most undifferentiated HSCs exist in several states closely related in UMAP and FDL space, but with distinct transcriptomes (**Fig. 2A, fig. S3, fig. S8A**). The subsets HSC-1, HSC-2, and HSC-5 were similar, enriched for ribosomal proteins and the important atypical homeodomain protein HOPX. Three subsets were particularly distinguishable: HSC-6 enriched for cell cycle genes, HSC-4 enriched for the histone acetyltransferase NAA40 and the D-type IL-17 receptor, and HSC-3. While most HSC states were present across all ages, HSC-3 was largely specific to prenatal development, increasing prenatally and becoming the dominant HSC state at 20 weeks’ gestation before diminishing postnatally; consistent with overall heterogeneity of fetal HSCs (**Fig. 4A**)(*27*). HSC-3 expressed several transcripts suggestive of either a uniquely activated state *in vivo*, or the effect of tissue processing (*28, 29*). We therefore sought to identify a surface marker for prospective isolation of HSC-3. Differential expression analysis of transcripts encoding cell surface proteins highlighted CD69 as specifically expressed in HSC-3 (**Fig. 4B**). Since CD69 is known primarily as a marker of activated T-cells, we corroborated the existence of CD69^+^ HSC populations in three additional mid-gestation fetal liver samples and independently published human FL datasets (**fig. S9**)(*30-33*).

**Figure 4.**
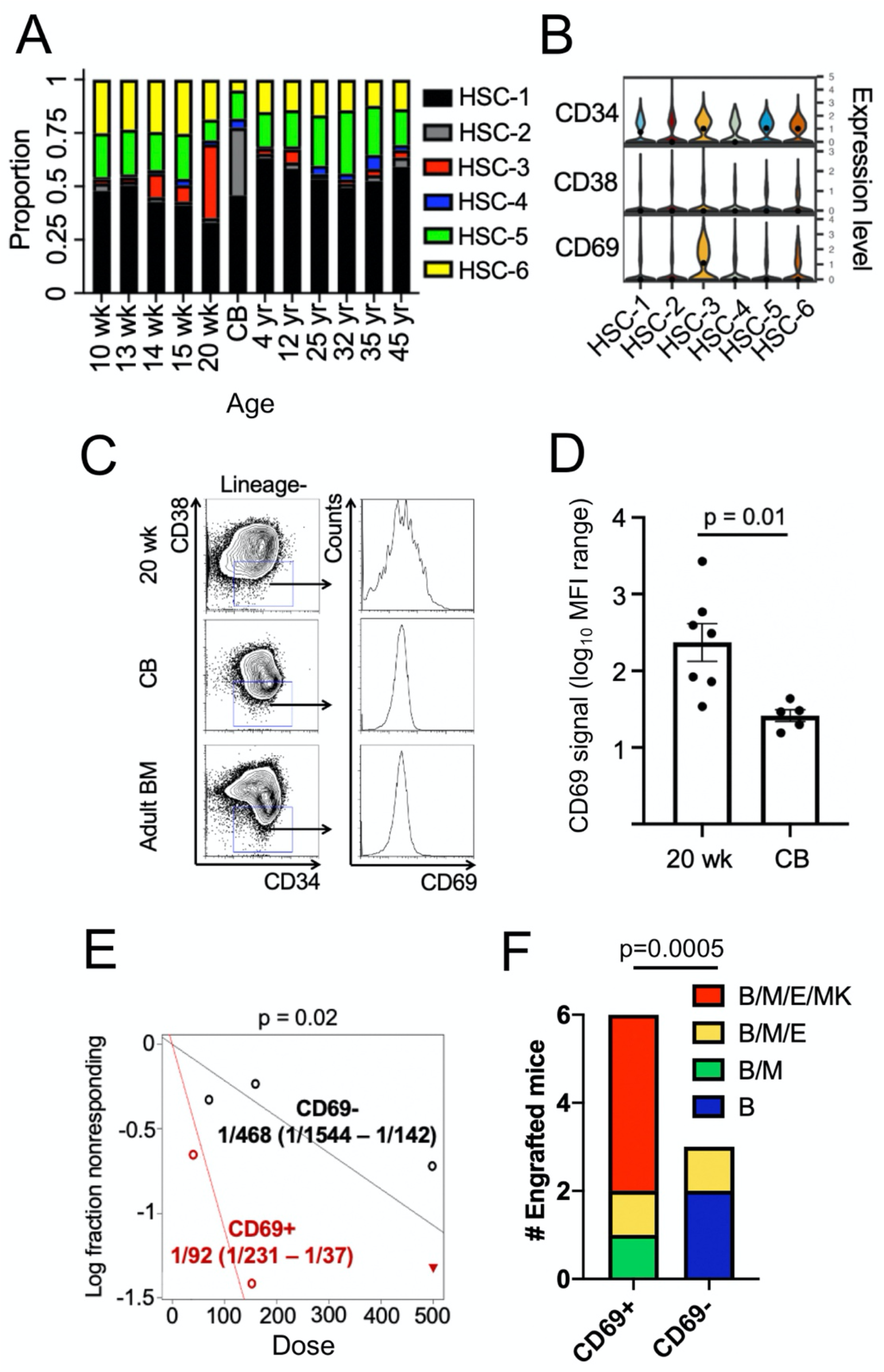
A transient HSC state in human fetal hematopoiesis. **(A)** HSC populations 1-6 at each age as a proportion of all HSC clusters. **(B)** Expression of transcripts encoding CD34, CD38, and CD69 in HSC1-6. **(C-D)** Following 48 hours of mitogenic stimulation, surface expression of CD69 was analyzed in the Lineage^-^CD34^+^CD38^-^ HSC/MPP population by flow cytometry, and the range of signal quantified. **(E-F)** CD69^+^ or CD69^-^ Lineage^-^CD34^+^CD38^-^ cells were transplanted into NSG recipients, and human chimerism and lineage outcome analyzed at 12 weeks (X^2^ = 5.17, 1 DF; B=B-cell, M=myeloid, E=erythroid, MK=megakaryocyte).

We next analyzed CD69 protein expression in fetal HSPCs. In freshly thawed human FL HSPCs, we did not detect appreciable CD69 expression in the undifferentiated Lineage^-^ CD34^+^CD38^-^ compartment, possibly indicating loss of cell surface CD69 during processing (**fig. S8B**). However, upon mitogenic stimulation, we observed heterogeneity in expression of cell surface CD69 specifically in mid-gestation FL HSCs/MPPs, consistent with the observed heterogeneity in *CD69* transcript expression (**Fig. 4C-D, fig. S8B**). To determine whether CD69 expression distinguishes functionally distinct HSC/MPP subpopulations, following mitogenic stimulation we sorted CD69^+^ and CD69^-^ gestational week 20 or later FL Lineage^-^CD34^+^CD38^-^ HSPCs (**fig. S8C**) onto MS5 stroma cells cultured with cytokines as well as in the presence or absence of erythropoietin at limiting dilution to quantify clonogenic potential and single cell lineage outcomes. Across multiple donors, mid-gestation CD69^+^ HSCs/MPPs were consistently more clonogenic with higher multilineage capacity (**fig. S10A-D**).

These findings indicate the CD69^+^ fraction of stimulated Lineage^-^CD34^+^CD38^-^ week 20 FL is enriched for HSCs compared to CD69^-^ cells. To test this, we performed *in vivo* limiting dilution xenotransplantation in NOD.Cg-*Prkdc*^*scid*^ *Il2rg*^*tm1Wj*^*/*SzJ (NSG) mice. Upon measuring human chimerism in recipient mouse bone marrow at twelve weeks following transplantation, we found CD69^+^ cells contained a significantly higher proportion of long-term engrafting cells compared to CD69^-^ cells (**Fig. 4E, fig. S10E**). Mice bearing CD69^+^ grafts showed a higher frequency of multilineage and particularly erythroid engraftment, compared to predominantly unilineage B-lymphoid production of CD69^-^ grafts (**Fig. 4F**). Lineage^-^CD34^+^CD38^-^CD69^+^ cells also reconstituted the Lineage^-^CD34^+^CD38^-^ compartment in recipient mice (**fig. S10F**). These findings uncover a mid-gestation human HSC state with potential for efficient multipotent engraftment.

Our findings show that during human development and aging, HSPCs temporally shift their transcriptional states to tailor mechanisms of lineage commitment and differentiation trajectories to support age-appropriate physiology. Although human HSPC ontogeny changes over time, HSCs and MPPs appear heterogeneous with skewed HSC fate probabilities and lineage biased MPPs at all ages. Within mid-gestation fetal liver, a transient HSC state emerges; engagement of this state may ensure balanced hematopoiesis prior to birth or efficiently populate the newly-formed bone marrow niche. Age-specific HSPC states can be retained or pathologically coopted in leukemia, and these states can shift from diagnosis to relapse as an apparent mechanism of therapeutic evasion. Our reconstruction of developmental hematopoiesis will serve as a foundation for future studies of human hematopoietic and immune development, aging, and age-specific blood disorders (*7*).

## Acknowledgments

We thank Harvey Lodish, Stuart Orkin, Caleb Weinreb, and Omer Yilmaz for helpful discussion.

## Author contributions

H.L. and R.G.R. designed the project and the experiments. H.L., V.M., D.W., and R.G.R. performed the experiments. J.E., A.A., L.G., S.Z., V.L.B., E.M., S.G., A.R., E.L.R., and G.S. developed analytical pipelines. H.L., J.E., A.A., L.G., S.Z., V.L.B., E.M., G.C., S.G., E.L.R., G.S., and R.G.R. analyzed the data. J.W., G.Q.D., and S.T.C. procured samples, funding, and reagents. All authors contributed to the writing and editing of the manuscript.

## Data and materials availability

Sequencing data from this study will be made available through the National Institutes of Health database of Genotypes and Phenotypes (dbGAP), for which a submission has been initiated. All other data is available in the main text or the supplementary materials.

## Supplementary Materials

Materials and Methods

Figure S1-10

Table S1

References (34-44)

