## Supplemental Files for "Hematopoiesis at single cell resolution spanning human development and maturation"

**This PDF file includes:**

Materials and Methods
Figs. S1 to S11

**Other Supplementary Materials for this manuscript include the following:**

Table S1

#### Materials and Methods

*Human cells and tissue*.
Human cord blood or adult bone marrow cells were purchased either from Stem Cell Technologies or AllCells. Human child bone marrow cells (age 4 or 12 years) were procured from healthy bone marrow donors enrolled on a clinical research protocol approved by the Human Subjects Protection Committee of the Dana-Farber/Harvard Cancer Center. Written informed consent was obtained before sample collection in accordance with the declaration of Helsinki. Human fetal liver cells were procured under the regulation of an approved Institutional Research Board protocol at both Boston Children's Hospital and The University of Pennsylvania.

*Fetal liver dissociation and cell isolation.*
Human fetal liver tissue was cut into 1mm diameter pieces using scissors and treated with 2 mg/mL Collagenase IV for 20 minutes at 37 degrees Celsius. Notably, all fetal liver specimens were cut into the same sizes and treated for the same duration in Collagenase IV given the known effects of differential collagenase treatment on gene expression in scRNA-seq experiments (*1*). Tissue was passed through a 70 micron filter and the resulting single cell suspension was subjected to Ficoll extraction. Mononuclear cells were magnetically enriched for CD34 using the Human CD34 Enrichment Kit (Miltenyi) and then cryopreserved in a solution of 80% fetal bovine serum and 20% dimethyl sulfoxide.

*Flow cytometry and cell sorting.*
Antibodies used in these studies included human lineage cocktail-Pacific Blue (CD3/14/1619/20/56), anti-human CD34 PE-Cy7 (8G12), anti-human CD38 PE (HB7), anti-human CD69 APC, anti-human CD235a PE-Cy7 (GA-R2), anti-human CD33 APC (P67-6), anti-human CD41a FITC, anti-human CD19 PE (HIB19), anti-human CD45 PE-Cy5 (all from BD Biosciences) and anti-mouse CD45.1 APC-Cy7 (clone A20, Biolegend). Flow cytometry analysis was performed on either a BD LSRII or LSR Fortessa. Cell sorting was performed on a BD FACS Aria.

*Cell culture.*
For mitogenic stimulation, human HSPCs were cultured in XVIVO medium (Lonza)supplemented with 1% bovine serum albumin, 1% penicillin, and 1% streptomycin (all from Gibco), 100 ng/ml recombinant human stem cell factor (SCF; R&D Systems), 100 ng/ml recombinant human FLT3-ligand (FLT3L; Peprotech), and 50 ng/ml recombinant human thrombopoietin (TPO; R&D Systems). For colony formation assays, 2000 cells were cultured in MethoCult H4434 Classic Methylcellulose Medium (Stem Cell Technologies) for 14 days at which time colony formation was quantified. For multilineage clonal assays with limiting dilution, cells were sorted at doses of 4, 2, or 1 cells into wells of 96 well plates seeded 48 hours previously with MS5 stromal cells (5000 cells/well) in Myelocult H5100 (Stem Cell Technologies) supplemented with 100 ng/ml SCF, 50 ng/ml TPO, 10 ng/ml FLT3L, and 25 ng/ml recombinant human interleukin-7 (IL-7; R&D Systems) with or without 2 units/ml human erythropoietin (EPO). Assays were fed weekly with fresh medium with cytokines. After 4 weeks, individual wells were visually scored for clonal outgrowths (blinded to the cell source) and single cell-derived outgrowths were picked for flow cytometry and morphologic analysis. The frequency of clonogenic cells was calculated using Extreme Limiting Dilution Analysis (*2*).

*Morphology.*
Human cells were spun onto slides and stained with May-Grunwald and Giemsa stains (Sigma) prior to morphologic analysis.

*Xenotransplantation.*
NOD.Cg-*Prkdc*$^{scid}$ *Il2rg*$^{tm1Wj}$/SzJ (NSG) mice were conditioned with 275 rad ionizing radiation prior to injection of the indicated cell numbers via the tail vein. 12 weeks following transplantation, human chimerism was quantified by flow cytometry, and engraftable stem cell frequency was calculated by limiting dilution analysis (*2*).

*InDrop Single-cell RNA-sequencing*
Cryopreserved fetal liver, cord blood, and bone marrow CD34-enriched cells were thawed and FACS-sorted for CD34+ cells. Sorted CD34+ cells were submitted to the Harvard University Single Cell Core for inDrop single cell RNA library preparation. Single cell RNA libraries were sequenced on an Illumina NextSeq 500. Raw sequencing reads were processed using the inDrop pipeline (https://github.com/indrops/indrops) using default parameters. The GRCh38 reference genome was used for alignment of sequencing reads.

*Single-cell RNA-Seq processing and cell-type clustering and analysis*
Across all samples, 64978 cells were called. Count matrices of genes x cells were imported in the R v. 3.6.0 statistical environment and Seurat v.3.0 was used as the primary analytical package (*3*). Count matrices were merged into a single Seurat object. Normalization to ten thousand transcript per cell barcodes was run on each sample individually, log-transformed, and variable features were identified using the vst method based on the top 2000 feature. The samples were then integrated using Seurat's IntegrateData procedure, using 2000 anchors and 30 dimensions (*4*). Dimensionality reduction using principal component analysis was performed and the top 30 principal components were used to build the kNN graph, considering 10 nearest neighbors. The resulting graph was partitioned using the Louvain shared nearest neighbor (SNN) modularity optimization-based clustering algorithm at resolution 0.8 to identify clusters. Iterative visualizations and plotting of QC metrics led to the exclusion of cells with low-transcript and high-mitochondrial content, as well as cells displaying endothelial and liver markers. The final object encompassed 14 datasets, resulting in 38873 cells and 41569 genes being retained for downstream analysis. Graph partition resulted in 31 distinct clusters. Cluster-specific gene markers were identified from the un-integrated, normalized gene expression data using a Wilcoxon test. Clusters were manually examined and merged based on their frequencies and gene expression profiles, leading to 21 clusters being retained for all downstream analyses. Proportions of cells from each sample and each cluster were tallied and represented as bar-charts.

Functional signatures were retrieved from the literature (*5-8*) and manually curated for cell/developmental-stage specific markers.
*Erythroid*- KLF1, CA1, HBB, HBD, GYPA, TFR2
*Megakaryocyte/Platelet*- GP1BB, PBX1, ITGA2B, PF4, VWF, PLEK
*Granulocyte*- SPI1, GFI1, CEBPA, ELANE, MPO, CSF1R, CTSG, PRTN3, AZU1
*Monocyte/Dendritic Cell*- IRF7, IRF8, CCR2, MPEG1, SPIB, IGKC, MS4A6A, ANXA2, MNDA, FCN1, CD14, CSAR1, CLEC4A, CLEC10A, FCER1A, CLEC4C, PTPRA, TCF4
*Basophil/Eosinophil/Mast Cell*- HDC, LMO4
*Lymphoid*- EBF1, ID3, DNTT, CD79A, VPREB1, RAG1, RAG2, MME, PAX5, CD19, CD79A, MS4A1, BANK1, MZB1, IGLL5, CD3D, CD3G, IL32, IL7R, TCF7, CCL5, GZMK, CD8A, KLRB1, KLRD1, NCAM1

Each signature was scored in each cell by summing transcript counts normalized per 10K transcripts across each gene present in the signature. Scores were visualized by projection over integrated UMAP embeddings.

To generate the HSC-specific heatmap, a new integrated Seurat object encompassing HSC-1 to -6 clusters was generated, normalized and cluster-specific markers were elicited using Wilcoxon and mast statistical frameworks (*9*). Markers that were significant by either or both statistics that also displayed a difference in percent positive cells > 30% between each cluster and the background were retained for plotting on the heat-map. In addition, the top 20 some markers that discriminated HSC-5 and -6 from the all-sample integrated object were added to the list. Cells from each cluster were down-sampled to 200 cells per cluster.

*Visualization of single-cell RNA-Seq data*
Using Seurat v.3.0 (*3*), the UMAP embedding was generated based on the same top 30 principal components of the integrated dataset (*10*). For the purpose of cell lineage reconstruction, the PAGA algorithm (Scanpy v. 1.4.4.post1, python3 v. 3.6.4 environment) (*11*) was run on cells present in the final Seurat integrated object, using log-transformed data and the same cluster calls. Principal-component analysis was re-run in the Scanpy/PAGA environment using arpack as a solver. A kNN graph was constructed based on 7 neighboring cells and the first 20 principal components. The underlying graph abstraction was learned using the tl.paga function and served to guide a force-directed network representation of the cells (using ForceAtlas2, as interfaced by PAGA) (*12*). Resulting cell coordinates were retrieved and used for plotting gene expression intensities and other metrics of interest in the Seurat environment.

*SingleCellNet analysis*
To quantitatively assess the molecular similarity between cell types and states identified in our dataset relative to leukemia samples generated previously (*7, 13*) we employed SingleCellNet to construct cell-type specific classifiers from our scRNA-seq data (*14*). These leukemia datasets were imported into CellRouter for filtering purposes (*15*). Briefly, for each B-ALL sample, we selected only cells with expression of CD19 higher than zero across the entire dataset. Then, we selected CD19+ cells by keeping only cells with an expression of CD19 higher than the mean expression. Then, we queried CD19+ cells from each patient sample through the SingleCellNet classifiers. For AML leukemia samples, we used metadata (cell type annotations classification of malignant or non-malignant cells) as originally published by the authors to classify malignant cells relative to the SingleCellNet models trained using our dataset.

*Computation of Fate Probabilities by PBA*
Population balance analysis (PBA) is a graph-based probabilistic methodology for inferring lineage trajectories from static snapshot data such as scRNA-seq (*16*). PBA relies on a number of assumptions to allow for a constrained, unique output: cell expression dynamics are Markovian, absence of rotational gene-expression dynamics, and large number of sampled cells. We apply PBA to each time point, 10 weeks to 45 years old, to assess how the lineage trajectories or fate biases change with time. The algorithm's primary inputs are the single cell expression matrix (cell x gene), lineage-specific "sink" matrix (cell x 6 lineages) and the diffusion constant. From the expression profile, a knn-graph is formed (n-neighbors set at 20), which is used to propagate probability mass between cell states. In this framework, we define "sinks" as the cells in the system that are the most terminally differentiated, where mass would be exiting the most. The lineage specific "sinks" are assigned based on enrichment for well-known, predefined markers for each of our 6 major lineages. The lineage "sink" matrix ultimately defines the fluxes or the net rate of loss associated with terminally differentiated cell states for each

lineage. Lastly, the diffusion constant defines the level of stochasticity cell states have in traversing other more distant cell states. We chose a diffusion constant of 0.2, as the recommended range from the PBA study was between 0.1-0.3, and we find this to produce the most stable results. Given the flux, the "compute_fate_probability" function computes the probability that in a random walk a given cell state will be absorbed by nodes in the graph associated to each of the terminal sinks.
The initial fluxes for the sinks are set at $1/D$, and we iteratively modify the fluxes such that difference between the summed output fate probabilities per lineage and the observed fate probabilities, as defined by pre-defined lineage-specific cluster annotations, are minimized (**fig. S11** showing expected vs actual).

*Optimal Transport-based computation of fate probabilities by StationaryOT*
Similarly to PBA, stationary optimal transport models biological processes in equilibrium as a diffusion-drift process subject to cell division and death (*17*). While these methods share a problem formulation, they take theoretically distinct approaches with stationary OT using entropy-regularised optimal transport to infer dynamics. As with PBA, the inputs to stationary OT include an expression matrix and assignments of sink cells by lineage, and the output is a cell-by-cell transition matrix from which fate probabilities may be computed. Unlike the flux rates which are not strictly enforced in the PBA method, the mass entering each sink in stationary OT can be exactly controlled through sinks weights and does not rely on ad-hoc flux tuning by the user in order for the overall fate probabilities to match the observed proportion of cells. The level of diffusion is controlled by the entropy regularisation parameter $\epsilon$, and estimates of cell growth rates are provided to model the effect of cell death and division.

Using stationary OT, we computed a transition matrix for each time point, assigning a total of 100 sinks proportional to the expected amount of terminal mass in each of the six lineages from a cluster-based annotation. For consistency, the same sinks were chosen as in the PBA analysis, and similar to PBA, we assumed no significant differences in growth rates in the different lineages. The couplings were calculated in a 50-dimensional PCA space based on the top 2000 highly variable genes. Based on empirical observations, we chose $\epsilon = \bar{C}$, the mean of the cost matrix.

A Python implementation of the stationary OT method is available at https://github.com/zsteve/statOT

*Elastic Net Gene Regression*
We perform elastic net regression to determine a set of genes ranked by their respective importance to lineage determination. We then visualize these genes and their rankings on a heat map with respect to the age of samples. Genes whose ranking is relatively constant over time are separated from those whose ranking varies, using an optimal transport based method.

Elastic net is a regression method that combines both $L_1$ and $L_2$ regularization (*18*). We apply the elastic net to gene expression matrices $E_t$, for each sampled age $t$. The regression is performed on a lineage-by-lineage basis against $f_{L,t}$, fate probabilities for a given lineage $L$, for cells from age $t$. In this context, the objective function for the elastic net is:

$$\frac{1}{2n}\left|f_{L,t} - E_t w\right|_2^2 + \alpha r_{L_1}|w|_1 + \alpha\left(1 - r_{L_1}\right)|w|_2^2$$

Where $n$ is the number of cells, $w$ is a vector of regression coefficients, $\alpha$ is a regularization coefficient, and $r_{L_1}$ is the ratio between the $L_1$ and $L_2$ terms. $r_{L_1}$ was fixed to be $10^{-3}$ as only a minimal degree of sparsity was desired. $\alpha$ was selected for each age and lineage by maximizing the $R^2$ of the fit. The Python package Scikit-Learn was used as the solver for the regressions (*19*).
The regression assigns a coefficient to each gene. The regression coefficient for a gene and a given lineage determines how much a unit change in expression (after normalization) of that gene affects a cell's probability of achieving that lineage. Although coefficients can be both positive and negative, for up-regulation favored genes and down-regulation favored genes, respectively, we will only consider those that are positive. We can then use the coefficient for each gene to determine its importance in lineage specification.

To visualize the outcome of the regressions, we created heat maps that show the relative coefficients of genes across ages. In the heat maps, we divided genes into two categories: consistent and variable. Consistent genes were defined as those that have similar (non-zero) regression coefficients across most ages. Variable genes were defined as those that have non-zero regression coefficients of similar magnitude only in a certain period of development (fetal, child, or adult). These heat maps can be seen in figure 3B and supplementary table S1.
To create the heat maps, we first normalized the regression coefficients for each gene so that they would form a probability distribution over time. That is, for a given lineage and a gene $g$, for each time $t$, its normalized regression coefficient is:

$$c_{g,t} = \frac{w_{g,t}}{\sum_{t_i \in \{times\}} w_{g,t_i}}$$

Optimal transport is then applied on the normalized coefficients to separate genes into constant and variable. For the consistent genes, a target uniform distribution across all ages was created. Note that ages were taken to be consecutive integers for all optimal transport computation. Then, the Earth Mover Distance (EMD) under the square euclidean metric was computed using the Python Optimal Transport package (available at https://pythonot.github.io) between each gene and this uniform distribution (*20*). EMD is the minimum cost associated with solving the Monge problem where the cost function is a metric (*21*). Consistent genes were taken to be those below the $k\%$ quantile for this distance. Variable genes were selected in a similar way, with the target distribution instead falling only on a period of ages (fetal, child, or adult). Again, the bottom $k\%$ quantile from each period was taken. For the featured figures and tables, $k$ was taken to be $15\%$. Each group was then sorted in descending order by regression coefficient. For the figure panel 3B, a curated list of genes was selected from these groups, the majority of which appear in the top 30 genes in their respective group. For the supplementary tables, a fixed number of genes were selected from the top of each group. On the heat map, the variable genes are ordered by their expected time, while the consistent genes are ordered alphabetically. Here, expected time is given by $\sum_{t \in \{times\}} t c_{g,t}$, and can be interpreted as the average time at which the gene has a lineage determining role. Finally, genes are colored at each time relative to the size of their own normalized coefficients. As a result of this procedure, the heat maps show genes that may have a lineage determining role at various periods of development.

*Data availability*
The sequencing data from this study will be made available through the National Institutes of Health database of Genotypes and Phenotypes (dbGAP), for which a submission has been initiated and will be accessible prior to publication

**Figure S1**

**A**

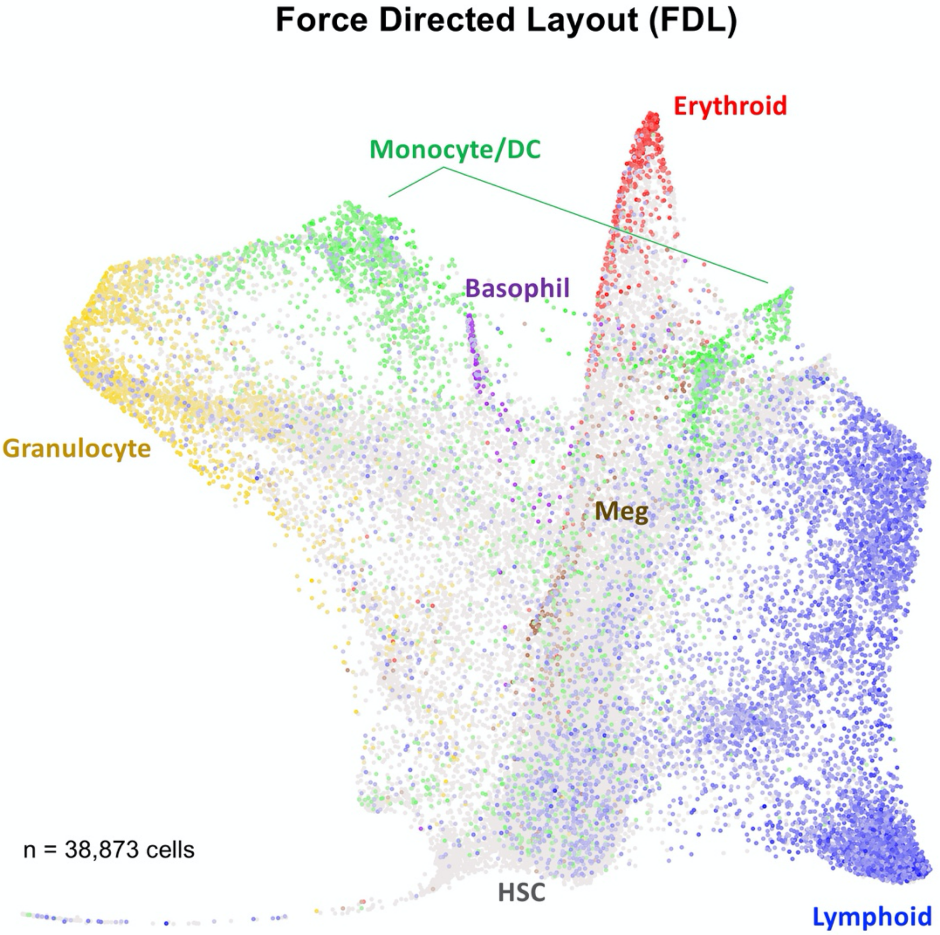
Force Directed Layout (FDL)
Erythroid
Monocyte/DC
Basophil
Granulocyte
Meg
n = 38,873 cells
HSC
Lymphoid

**B**

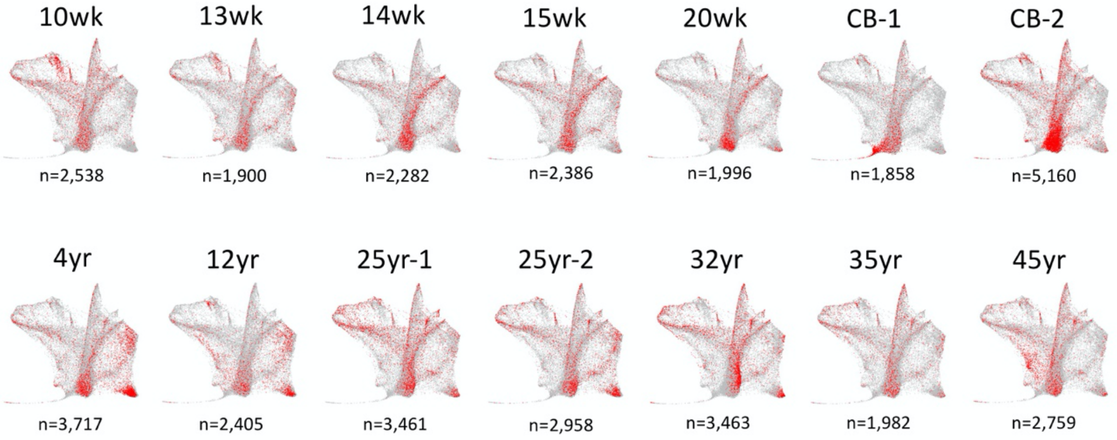
10wk
13wk
14wk
15wk
20wk
CB-1
CB-2
n=2,538
n=1,900
n=2,282
n=2,386
n=1,996
n=1,858
n=5,160
4yr
12yr
25yr-1
25yr-2
32yr
35yr
45yr
n=3,717
n=2,405
n=3,461
n=2,958
n=3,463
n=1,982
n=2,759

**Figure S1. Single cell transcriptome visualization via Force-Directed Layout (FDL). (A)** PAGA-abstracted Force-directed layout (FDL) of all single-cell transcriptomes in the study. **(B)** Distribution of individual cells for each sample displayed on FDL. Red dots indicate cells from that sample. Gray background dots indicate cells from all other samples.

**Figure S2**

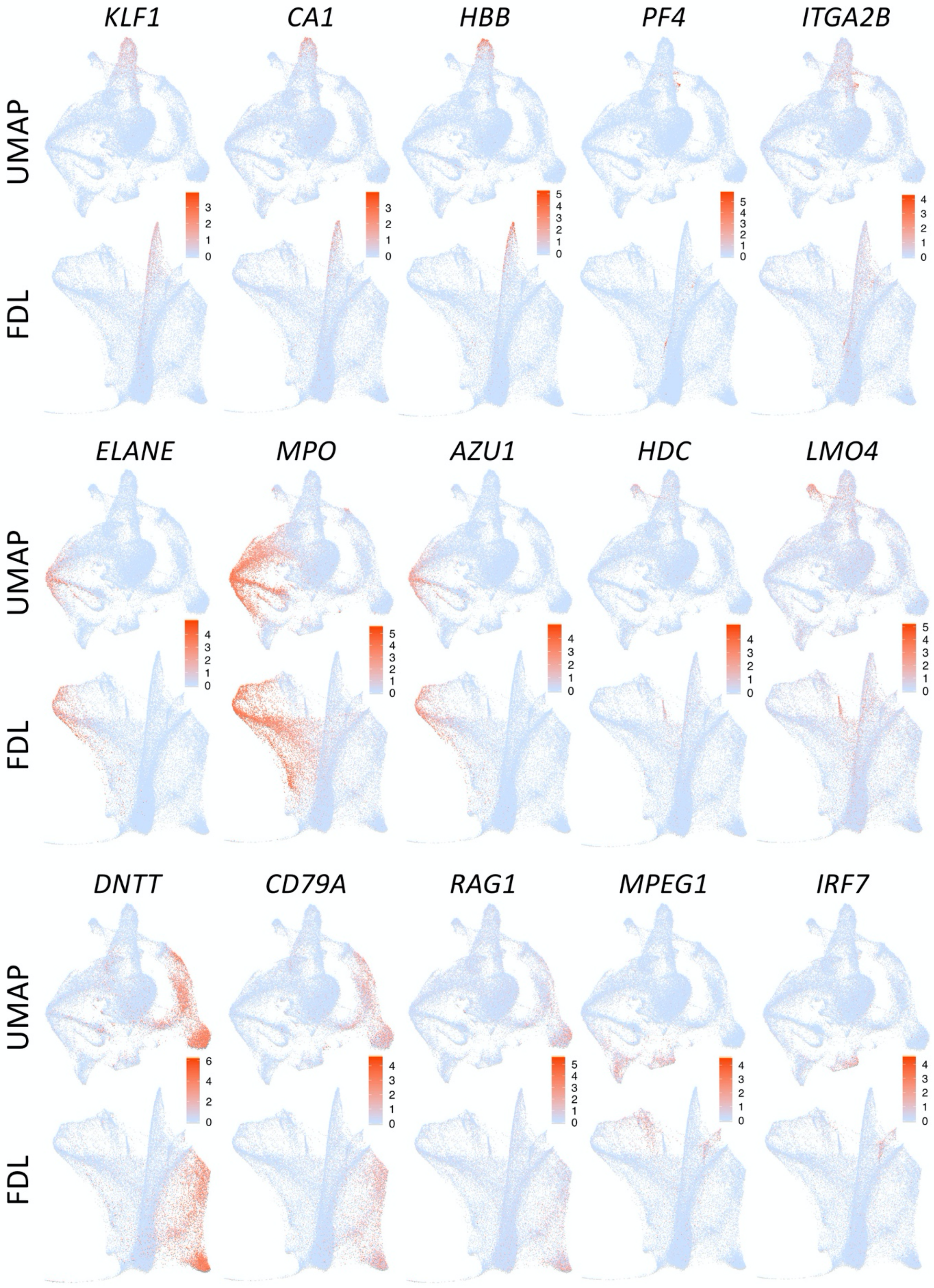
KLF1
CA1
HBB
PF4
ITGA2B
ELANE
MPO
AZU1
HDC
LMO4
DNTT
CD79A
RAG1
MPEG1
IRF7
UMAP
FDL

**Figure S2. Lineage-specific marker gene expression.** Relative expression of characteristic marker genes of the erythroid (KLF1, CA1, HBB), megakaryocyte (PF4, ITGA2B), granulocyte (ELANE, MPO, AZU1), basophil (HDC, LMO4), lymphoid (DNTT, CD79A, RAG1), and monocyte (MPEG1, IRF7) lineages overlaid on both UMAP and Force-Directed Layout (FDL) projection.

**Figure S3**

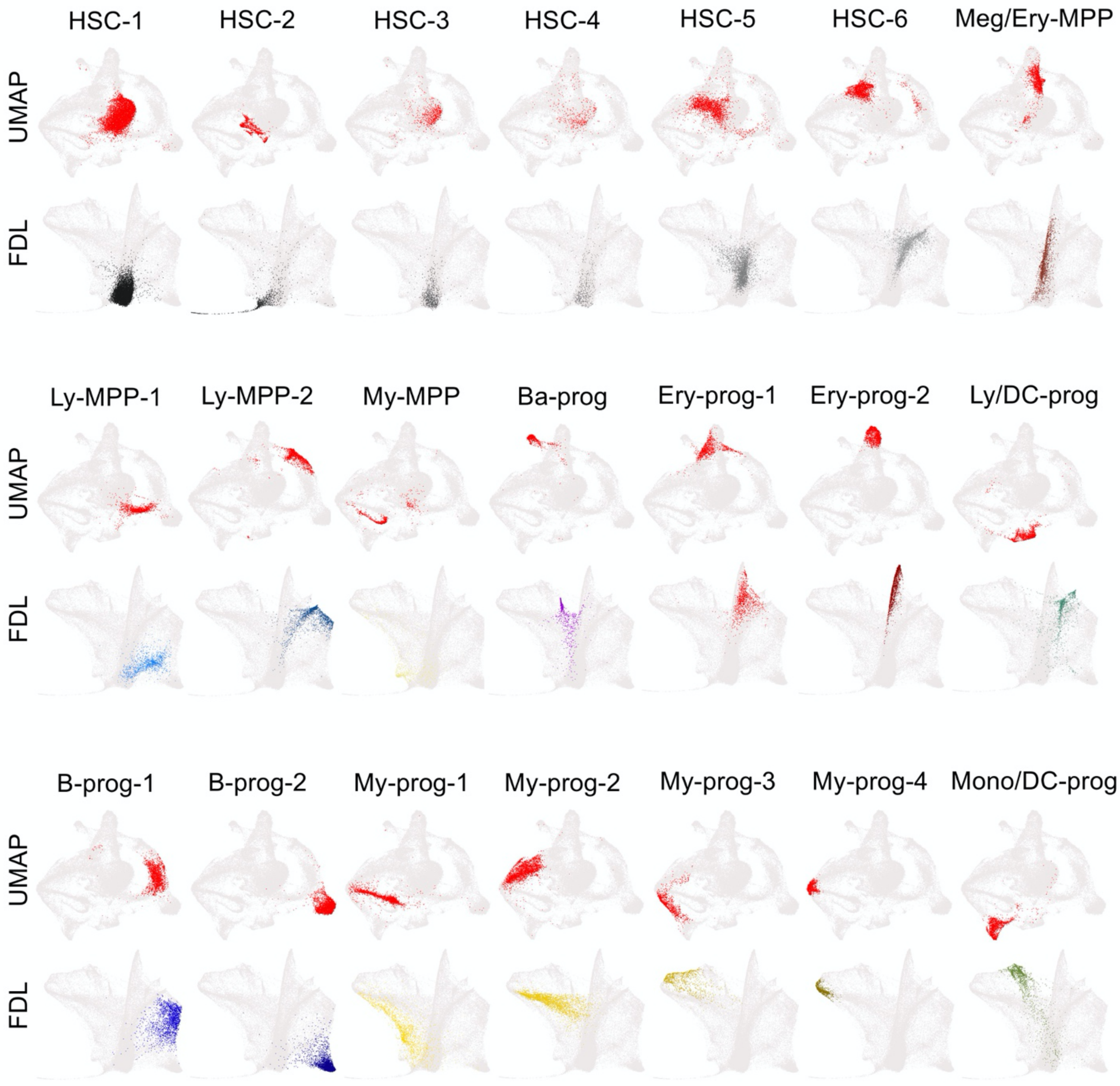
HSC-1
HSC-2
HSC-3
HSC-4
HSC-5
HSC-6
Meg/Ery-MPP
UMAP
FDL
Ly-MPP-1
Ly-MPP-2
My-MPP
Ba-prog
Ery-prog-1
Ery-prog-2
Ly/DC-prog
UMAP
FDL
B-prog-1
B-prog-2
My-prog-1
My-prog-2
My-prog-3
My-prog-4
Mono/DC-prog
UMAP
FDL

**Figure S3. HSPC subtype distribution in dimensionality reduction space.** Distribution of individual clusters from Louvain clustering on UMAP and Force-Directed Layout (FDL) embedding. For UMAP plots, individual clusters are colored in red, with all other clusters as background gray. For FDL plots, individual clusters are colored with the color scheme from Fig. 2, with all other clusters as background gray.

Figure S4

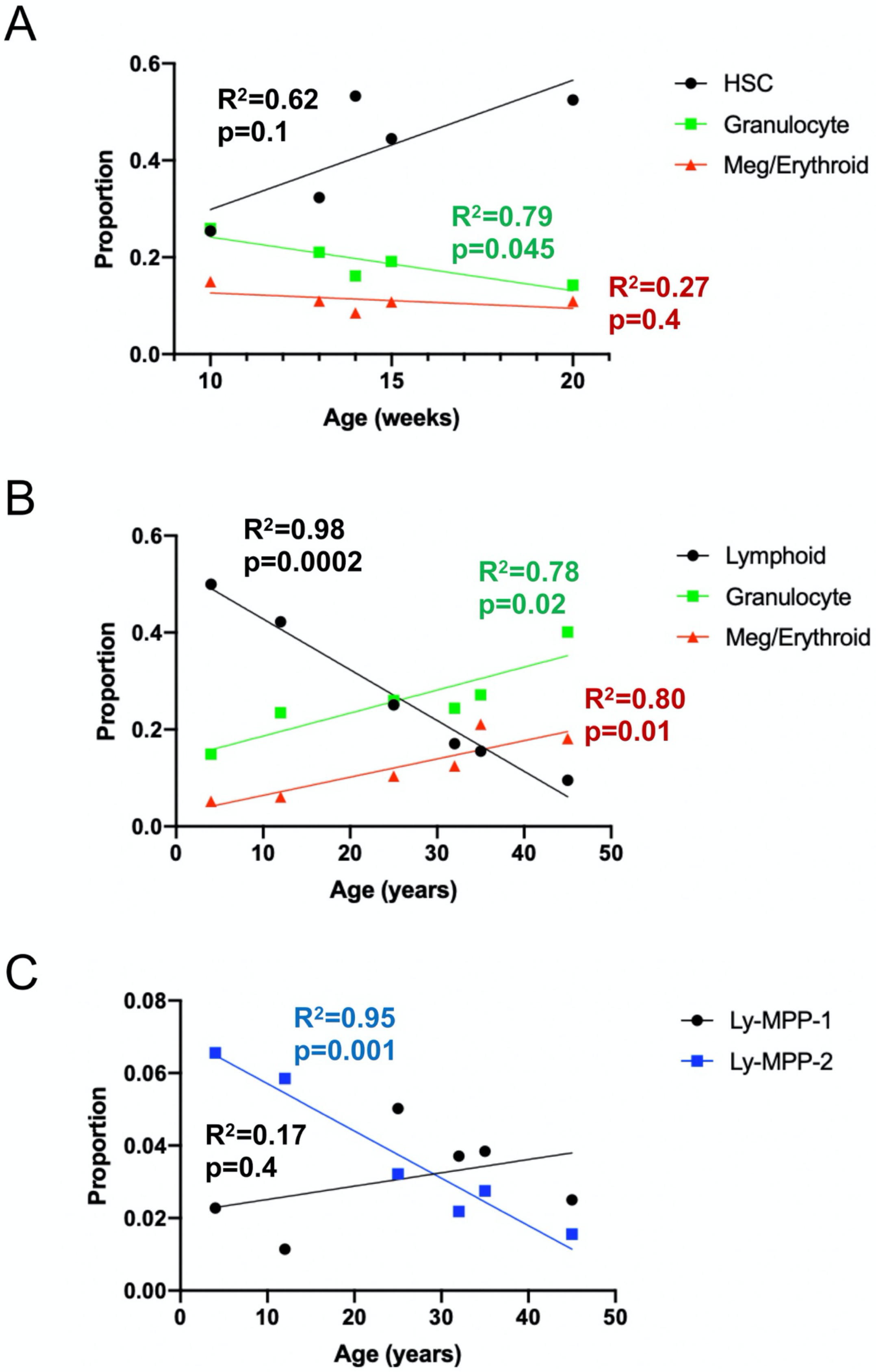

A
$R^2$=0.62
p=0.1
$R^2$=0.79
p=0.045
$R^2$=0.27
p=0.4
HSC
Granulocyte
Meg/Erythroid
Proportion
Age (weeks)
B
$R^2$=0.98
p=0.0002
$R^2$=0.78
p=0.02
$R^2$=0.80
p=0.01
Lymphoid
Granulocyte
Meg/Erythroid
Proportion
Age (years)
C
$R^2$=0.95
p=0.001
$R^2$=0.17
p=0.4
Ly-MPP-1
Ly-MPP-2
Proportion
Age (years)

**Figure S4. Change in lineage outputs with time. (A)** HSPC states assigned to the specific lineages were plotted over time as a proportion of all HSPCs at each age in prenatal specimens. **(B)** HSPC states assigned to the specific lineages were plotted over time as a proportion of all HSPCs at each age in postnatal specimens. **(C)** Ly-MPP-1 and Ly-MPP2 states as proportions of all HSPCs were plotted over postnatal time points. In all panels, changes over time were analyzed by simple linear regression with $R^2$ and *P* values shown.

**Figure S5**

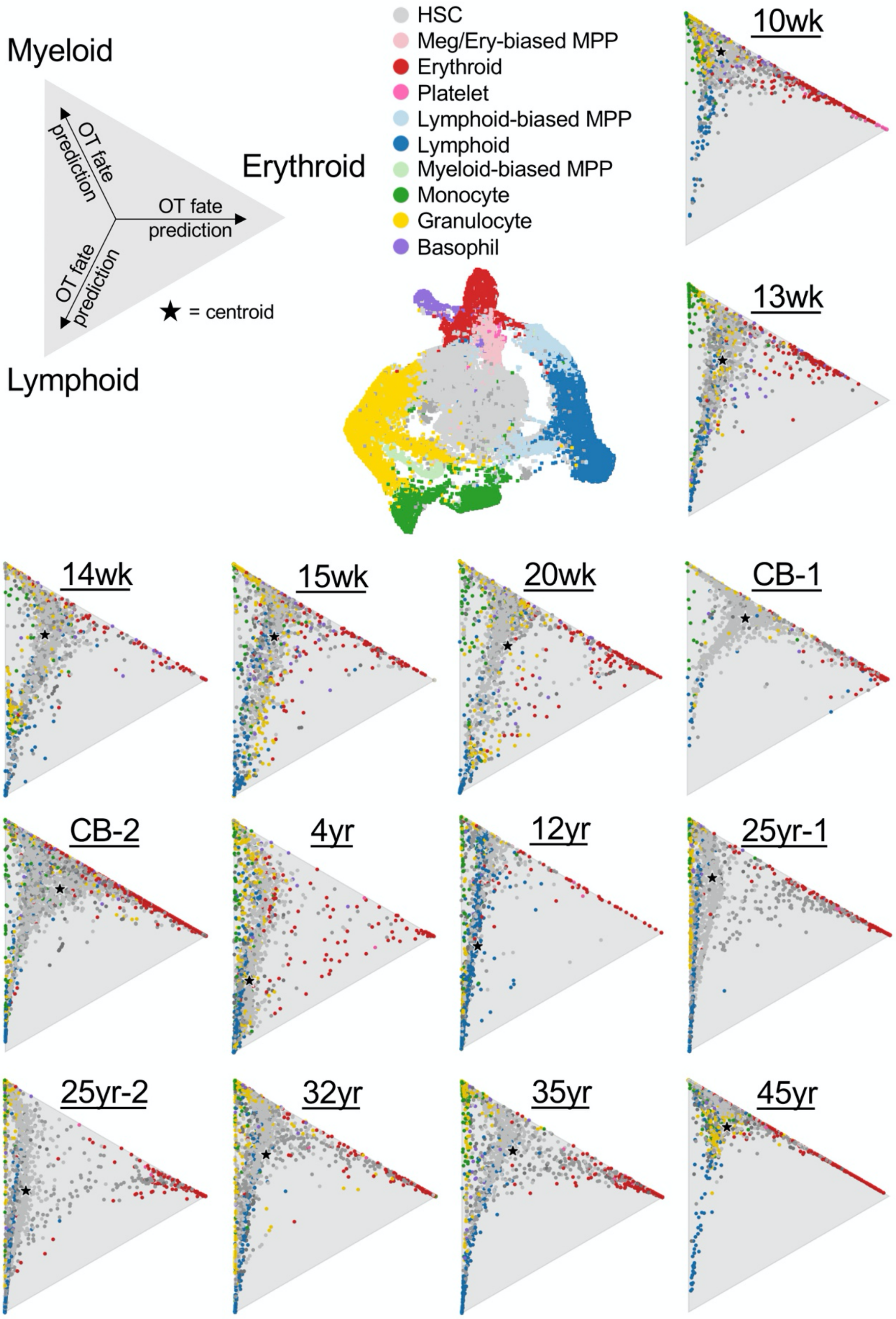

Myeloid
Erythroid
Lymphoid
OT fate prediction
OT fate prediction
OT fate prediction
★ = centroid
HSC
Meg/Ery-biased MPP
Erythroid
Platelet
Lymphoid-biased MPP
Lymphoid
Myeloid-biased MPP
Monocyte
Granulocyte
Basophil
10wk
13wk
14wk
15wk
20wk
CB-1
CB-2
4yr
12yr
25yr-1
25yr-2
32yr
35yr
45yr

**Figure S5. Optimal Transport lineage fate analysis.** Individual HSC fate probability of erythroid, myeloid, and lymphoid differentiation based on Optimal Transport (OT). Relative proximity of each cell dot to each vertex of the triangle indicates likelihood each cell differentiates towards each lineage. Centroid indicates average fate of all cells in that sample.

### Figure S6

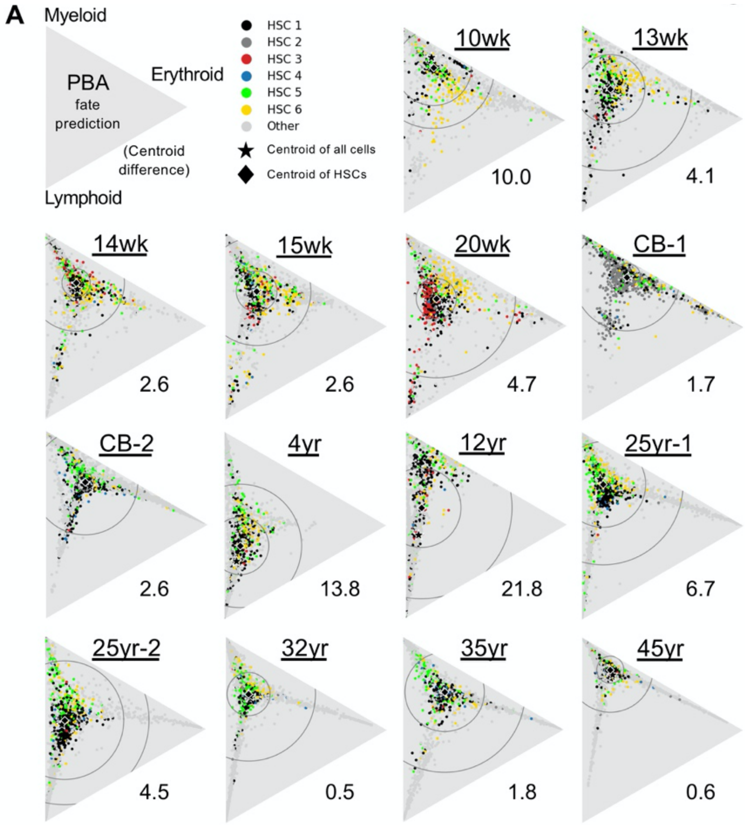
A
Myeloid
Erythroid
Lymphoid
PBA
fate
prediction
(Centroid
difference)
HSC 1
HSC 2
HSC 3
HSC 4
HSC 5
HSC 6
Other
Centroid of all cells
Centroid of HSCs
10wk
10.0
13wk
4.1
14wk
2.6
15wk
2.6
20wk
4.7
CB-1
1.7
CB-2
2.6
4yr
13.8
12yr
21.8
25yr-1
6.7
25yr-2
4.5
32yr
0.5
35yr
1.8
45yr
0.6

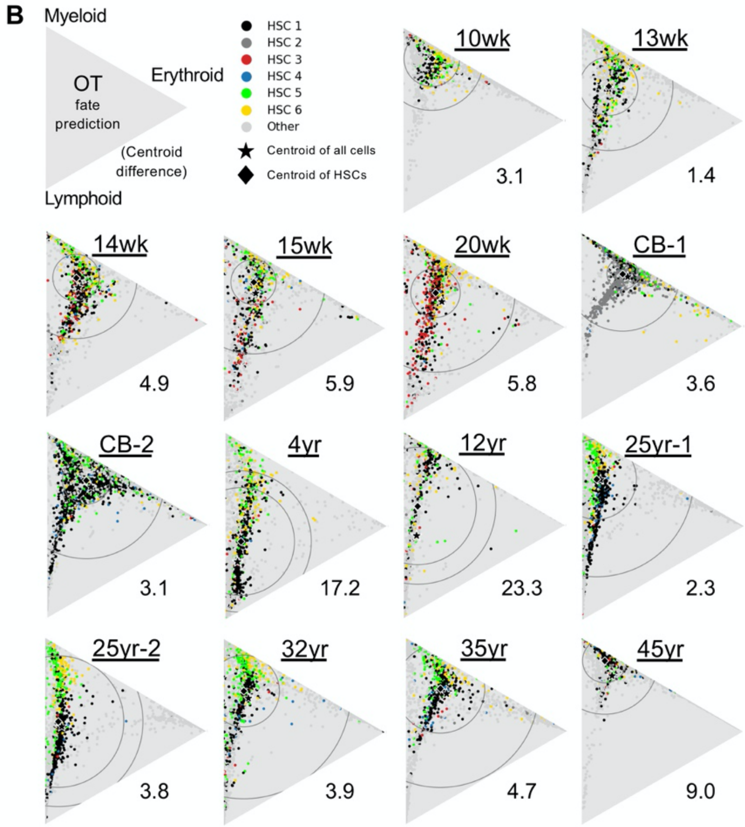
B
Myeloid
Erythroid
Lymphoid
OT
fate
prediction
(Centroid
difference)
HSC 1
HSC 2
HSC 3
HSC 4
HSC 5
HSC 6
Other
Centroid of all cells
Centroid of HSCs
10wk
3.1
13wk
1.4
14wk
4.9
15wk
5.9
20wk
5.8
CB-1
3.6
CB-2
3.1
4yr
17.2
12yr
23.3
25yr-1
2.3
25yr-2
3.8
32yr
3.9
35yr
4.7
45yr
9.0

**Figure S6. HSC lineage fate analysis by PBA and OT.** Individual cell fate probability of erythroid, myeloid, and lymphoid differentiation of HSC states based on **(A)** PBA, or **(B**) OT. Relative proximity of each cell dot to each vertex of the triangle indicates likelihood each cell differentiates towards each lineage. Centroids indicate average fate of either all cells in that sample, or only HSCs. Centroid difference is the discordance (distance) between the centroid for all cells and the centroid for only HSCs. Circles indicate 50% (inner) and 95% (outer) confidence intervals for all HSC fates.

**Figure S7**

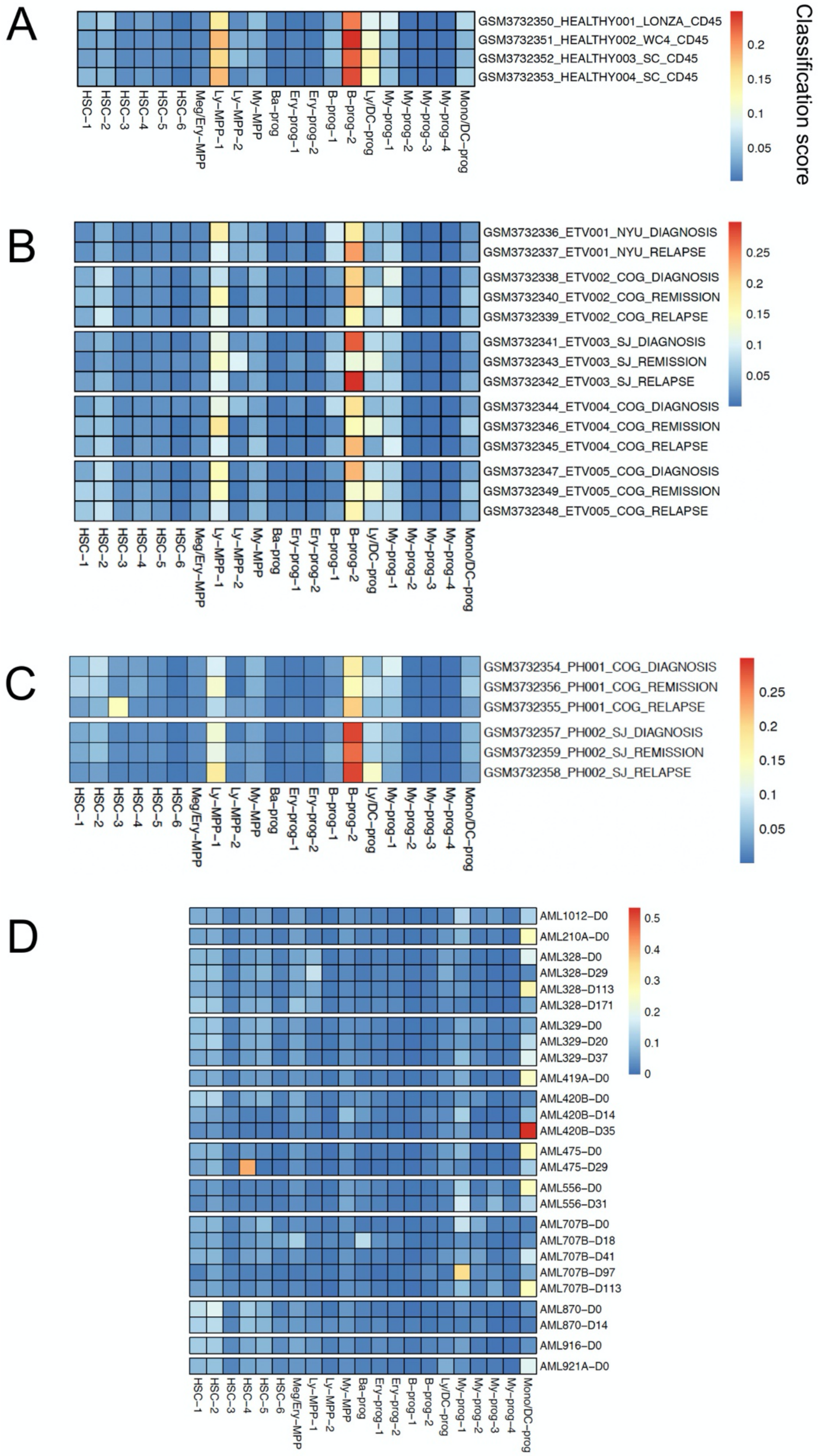
A
GSM3732350_HEALTHY001_LONZA_CD45
GSM3732351_HEALTHY002_WC4_CD45
GSM3732352_HEALTHY003_SC_CD45
GSM3732353_HEALTHY004_SC_CD45
Classification score
0.2
0.15
0.1
0.05
HSC−1
HSC−2
HSC−3
HSC−4
HSC−5
HSC−6
Meg/Ery−MPP
Ly−MPP−1
Ly−MPP−2
My−MPP
Ba−prog
Ery−prog−1
Ery−prog−2
B−prog−1
B−prog−2
Ly/DC−prog
My−prog−1
My−prog−2
My−prog−3
My−prog−4
Mono/DC−prog
B
GSM3732336_ETV001_NYU_DIAGNOSIS
GSM3732337_ETV001_NYU_RELAPSE
GSM3732338_ETV002_COG_DIAGNOSIS
GSM3732340_ETV002_COG_REMISSION
GSM3732339_ETV002_COG_RELAPSE
GSM3732341_ETV003_SJ_DIAGNOSIS
GSM3732343_ETV003_SJ_REMISSION
GSM3732342_ETV003_SJ_RELAPSE
GSM3732344_ETV004_COG_DIAGNOSIS
GSM3732346_ETV004_COG_REMISSION
GSM3732345_ETV004_COG_RELAPSE
GSM3732347_ETV005_COG_DIAGNOSIS
GSM3732349_ETV005_COG_REMISSION
GSM3732348_ETV005_COG_RELAPSE
0.25
0.2
0.15
0.1
0.05
C
GSM3732354_PH001_COG_DIAGNOSIS
GSM3732356_PH001_COG_REMISSION
GSM3732355_PH001_COG_RELAPSE
GSM3732357_PH002_SJ_DIAGNOSIS
GSM3732359_PH002_SJ_REMISSION
GSM3732358_PH002_SJ_RELAPSE
0.25
0.2
0.15
0.1
0.05
D
AML1012−D0
AML210A−D0
AML328−D0
AML328−D29
AML328−D113
AML328−D171
AML329−D0
AML329−D20
AML329−D37
AML419A−D0
AML420B−D0
AML420B−D14
AML420B−D35
AML475−D0
AML475−D29
AML556−D0
AML556−D31
AML707B−D0
AML707B−D18
AML707B−D41
AML707B−D97
AML707B−D113
AML870−D0
AML870−D14
AML916−D0
AML921A−D0
0.5
0.4
0.3
0.2
0.1
0

**Figure S7. Classification of human leukemia.** Human leukemia is typically classified on the basis of oncogenic mutations/translocations and differentiation lineage, with many types of leukemia biased toward certain ages. We asked whether our dataset could be utilized as a resource for assigning differentiation and maturation state of human leukemias. Using singleCellNet, we classified the CD19+ fraction of seven patient specimens variably comprising diagnosis, remission, and relapse of childhood B-acute lymphoblastic leukemia (B-ALL) of the *ETV6-RUNX1* (ETV) or *BCR-ABL* (PH) subtypes (*13*). **(A)** Healthy control CD19+ cells classified as Ly-MPP-1, B-prog-2 and Ly-derived DC-prog, the predominant postnatal lymphoid progenitors. **(B-C)** Active *ETV6-RUNX1* leukemias showed Ly-MPP-1 and B-prog-2 classification with loss of normal Ly/DC-prog populations, although in four of five *ETV6-RUNX1* cases and one of the *BCR-ABL* cases, classification shifted toward the more differentiated B-prog-2 state with relapse, suggestive of a mechanism of therapeutic evasion. At diagnosis, the first *BCR-ABL* case classified predominantly as Ly-MPP-1 and B-prog-2, but some cells classified as My-prog-1, suggestive of a hybrid myeloid identity. Interestingly, at relapse, cells classified as the fetal-specific HSC-3, suggestive of recruitment of fetal stem cell programs. Like healthy control CD19+ cells (in panel **(A)**, remission CD19+ cells generally classified as Ly-MPP-1, B-prog-2 and Ly-derived DC-prog **(D)** We classified acute myeloid leukemia (AML) scRNAseq datasets against our atlas (*7*). We found that AMLs generally segregated into two groups: those classifying as My-prog-1/Mono/DC-prog (e.g. 210A, 419A, 475, 556, 329) and those showing a more undifferentiated state, with these classifications correlating with morphologic differentiation reported previously (Figure 3D)(*5*). For those AMLs classifying as undifferentiated and for which post-treatment specimens were available, we observed acquisition of My-prog-1 and Mono/DC-prog classification post-treatment. Together, these data demonstrate that our atlas of human HSPC maturation can be used to reconstruct leukemic ontogeny, precisely define differentiation state, and uncover oncofetal programs.

Figure S8

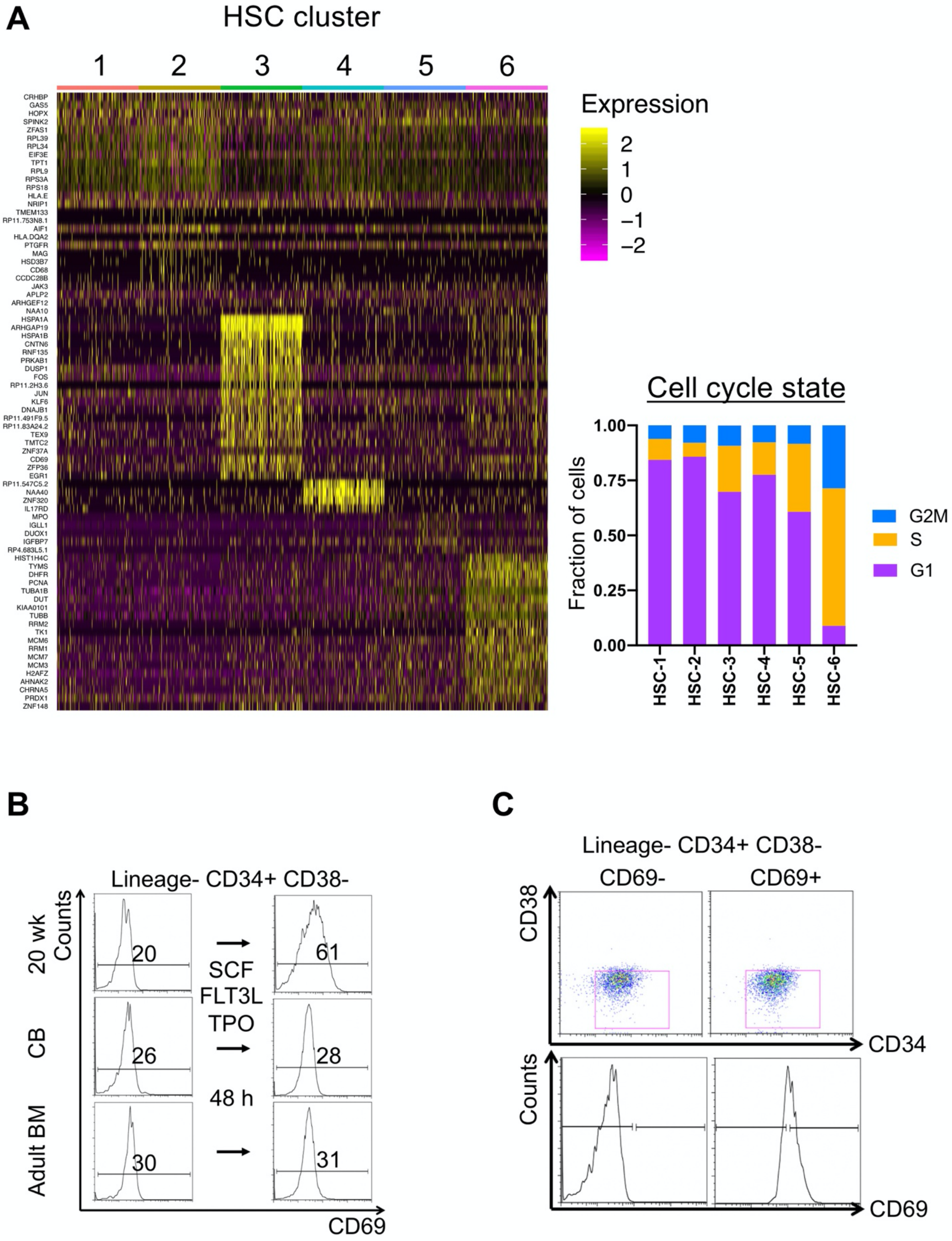
A
HSC cluster
1
2
3
4
5
6
CRHBP
GAS5
HOPX
SPINK2
ZFAS1
RPL39
RPL34
EIF3E
TPT1
RPL9
RPS3A
RPS18
HLA.E
NRIP1
TMEM133
RP11.753N8.1
AIF1
HLA.DQA2
PTGFR
MAG
HSD3B7
CD68
CCDC28B
JAK3
APLP2
ARHGEF12
NAA10
HSPA1A
ARHGAP19
HSPA1B
CNTN6
RNF135
PRKAB1
DUSP1
FOS
RP11.2H3.6
JUN
KLF6
DNAJB1
RP11.491F9.5
RP11.83A24.2
TEX9
TMTC2
ZNF37A
CD69
ZFP36
EGR1
RP11.547C5.2
NAA40
ZNF320
IL17RD
MPO
IGLL1
DUOX1
IGFBP7
RP4.683L5.1
HIST1H4C
TYMS
DHFR
PCNA
TUBA1B
DUT
KIAA0101
TUBB
RRM2
TK1
MCM6
RRM1
MCM7
MCM3
H2AFZ
AHNAK2
CHRNA5
PRDX1
ZNF148
Expression
2
1
0
−1
−2
Cell cycle state
Fraction of cells
1.00
0.75
0.50
0.25
0.00
HSC-1
HSC-2
HSC-3
HSC-4
HSC-5
HSC-6
G2M
S
G1
B
Lineage- CD34+ CD38-
20 wk
CB
Adult BM
Counts
20
61
26
28
30
31
SCF
FLT3L
TPO
48 h
CD69
C
Lineage- CD34+ CD38-
CD69-
CD69+
CD38
CD34
Counts
CD69

**Figure S8. Heterogeneity in HSCs and characterization and purification of HSC-3. (A)** Heatmap showing transcriptional signatures defining each of the indicated HSC populations and barplot of cell cycle distribution for HSC populations. **(B)** $CD34^+$ HSPCs from the indicated ages were analyzed immediately post-thaw (frozen following isolate from hematopoietic tissues) or following 48 hours of culture in the presence of mitogens (stem cell factor (SCF), Flt-3 ligand (FLT3L), or thrombopoietin (TPO) for the indicated markers, with representative results shown. Mean fluorescence intensities are indicated. **(C)** Representative results of post-sort purity analysis of sorted $Lineage^-$ $CD34^+$ $CD38^-$ $CD69^+$ or $CD69^-$ HSPCs prior to use in downstream clonogenesis or transplantation experiments.

**Figure S9**

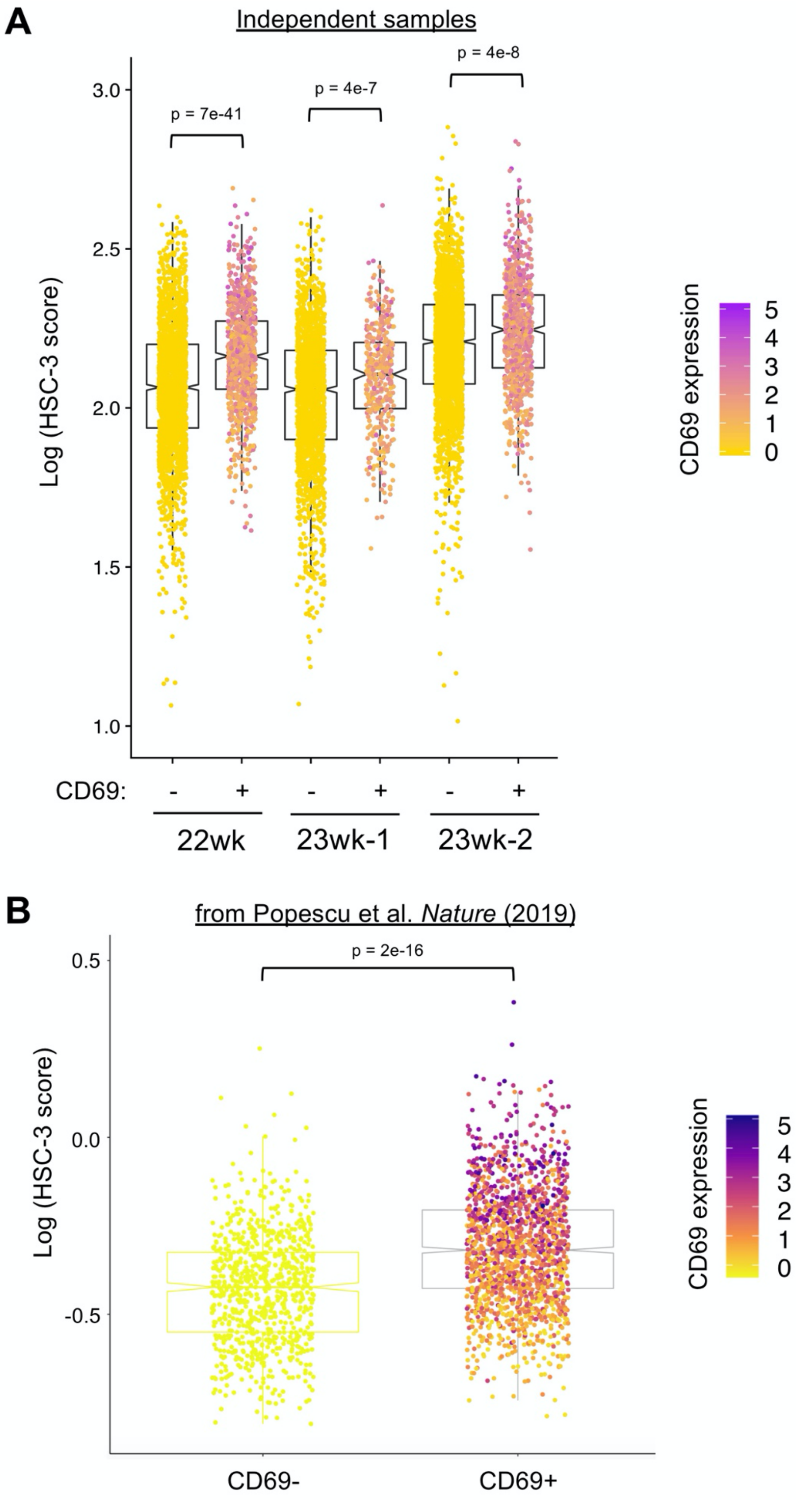
A
Independent samples
p = 7e-41
p = 4e-7
p = 4e-8
3.0
2.5
2.0
1.5
1.0
Log (HSC-3 score)
CD69 expression
5
4
3
2
1
0
CD69:
-
+
-
+
-
+
22wk
23wk-1
23wk-2
B
from Popescu et al. *Nature* (2019)
p = 2e-16
0.5
0.0
-0.5
Log (HSC-3 score)
CD69 expression
5
4
3
2
1
0
CD69-
CD69+

**Figure S9. CD69 expression marks HSC-3 identity. (A)** Three additional mid-gestation human fetal liver samples, independent from the fourteen samples analyzed in the study dataset profiled in Fig. 1, were profiled with single cell transcriptome analysis using the InDrop platform. Each sample was then classified as positive or negative for CD69 expression and similarity to HSC-3 cells measured using an HSC-3 cluster gene signature based on unbiased marker gene selection, with the log-transformed score value for each cell plotted. Color shading of each cell represents relative expression level of CD69. **(B)** CD34+ human fetal liver single cell transcriptomes profiled in Popescu, et al. 2019 (*22*) were analyzed in the same fashion, plotting log-transformed HSC-3 score value for CD69 positive and negative cells, with color shading of each cell represents relative expression level of CD69.

Figure S10

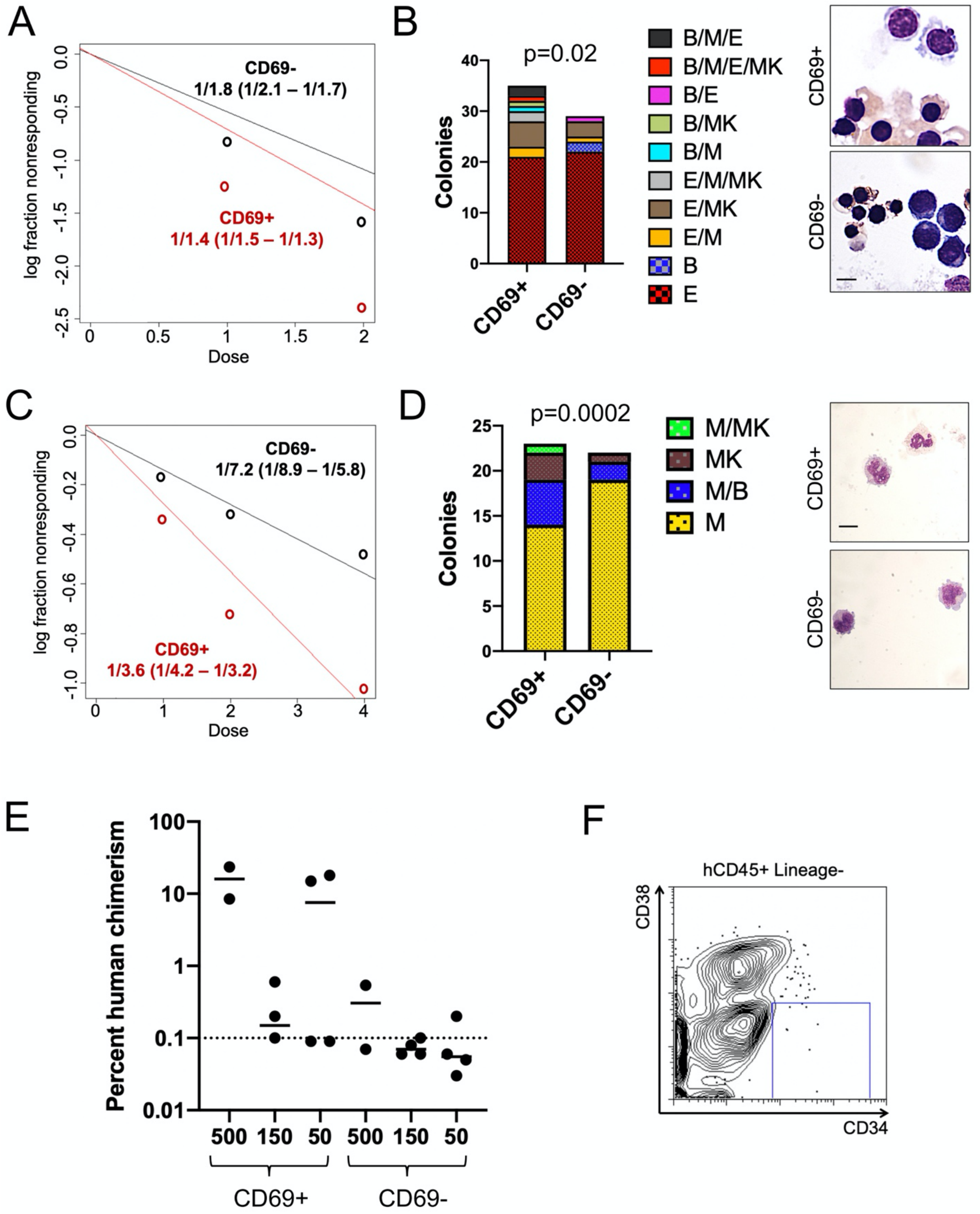

A
CD69-
1/1.8 (1/2.1 – 1/1.7)
CD69+
1/1.4 (1/1.5 – 1/1.3)
log fraction nonresponding
Dose
B
p=0.02
Colonies
CD69+
CD69-
B/M/E
B/M/E/MK
B/E
B/MK
B/M
E/M/MK
E/MK
E/M
B
E
C
CD69-
1/7.2 (1/8.9 – 1/5.8)
CD69+
1/3.6 (1/4.2 – 1/3.2)
log fraction nonresponding
Dose
D
p=0.0002
Colonies
M/MK
MK
M/B
M
E
Percent human chimerism
500 150 50 500 150 50
CD69+
CD69-
F
hCD45+ Lineage-
CD38
CD34

**Figure S10. CD69 expression defines a functionally distinct HSPC population. (A)** Lineage$^-$ CD34$^+$ CD38$^-$ CD69$^+$ or CD69$^-$ cells were sorted onto MS5 stroma with cytokines (recombinant human SCF, TPO, FLT3L, IL7, and EPO) at 4, 2, or 1 cells per well. After 4 weeks, colony outgrowths were scored, and clonogenic stem cell frequency calculated by limiting dilution analysis. Results pooled over two independent experiments from different donors. **(B)** Single cell-derived colonies were picked and lineage outcomes analyzed by flow cytometry, with representative morphology shown (Comparing unilineage versus multilineage colony outcomes: $X^2$ = 3.935, 1 df, p = 0.047. In image panels scale = 10 µm). **(C)** Lineage$^-$ CD34$^+$ CD38$^-$ CD69$^+$ or CD69$^-$ cells were sorted onto MS5 stroma with cytokines (recombinant human SCF, TPO, FLT3L, IL7 and without EPO in these experiments) at 4, 2, or 1 cells per well. After 4 weeks, colony outgrowths were scored, and clonogenic stem cell frequency calculated by limiting dilution analysis. Results pooled over two independent experiments from different donors. **(D)** Single cell-derived colonies were picked and lineage outcomes analyzed by flow cytometry, with representative morphology shown (Comparing unilineage versus multilineage colony outcomes: $X^2$ = 3.737, 1 df, p = 0.053. In image panels scale = 10 µm). **(E)** Percent human chimerism in each recipient mouse bearing the indicated dose of either Lineage$^-$ CD34$^+$ CD38$^-$ CD69$^+$ or CD69$^-$ cells at 12 weeks post-transplant. **(F)** Representative results of week 12 engraftment showing for markers of early HSPCs.

**Figure S11**

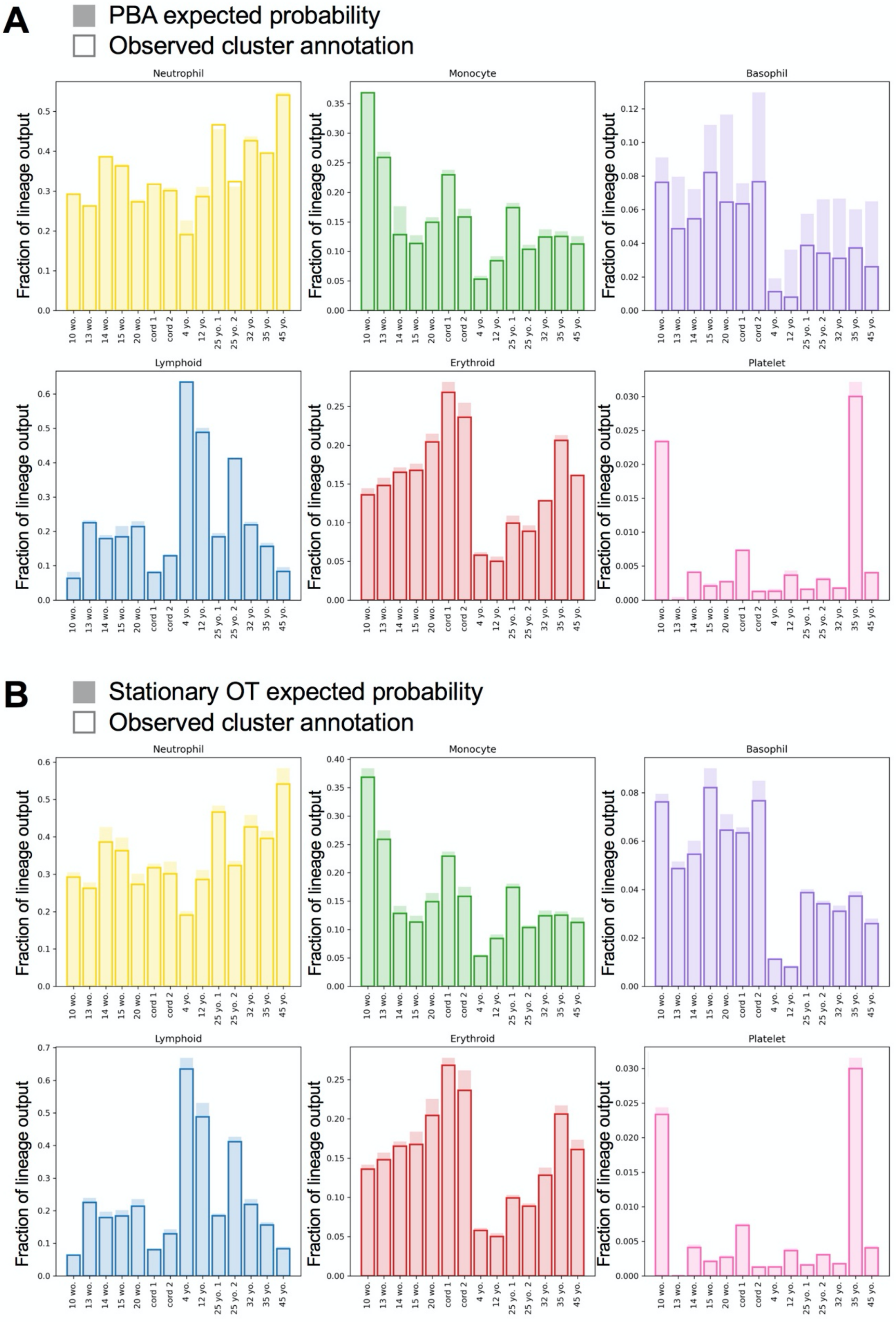
A
PBA expected probability
Observed cluster annotation
Neutrophil
Monocyte
Basophil
Lymphoid
Erythroid
Platelet
Fraction of lineage output
10 wo.
13 wo.
14 wo.
15 wo.
20 wo.
cord 1
cord 2
4 yo.
12 yo.
25 yo. 1
25 yo. 2
32 yo.
35 yo.
45 yo.
B
Stationary OT expected probability
Observed cluster annotation
Neutrophil
Monocyte
Basophil
Lymphoid
Erythroid
Platelet
Fraction of lineage output

**Figure S11. Comparison of Expected and Observed Probability Masses.** We compare the proportion of mass (as measured by fraction of total predicted hematopoietic output) by lineage and age in terminal fates for **(A)** PBA and **(B)** Stationary OT to the proportion of cells that reside in annotated clusters of differentiated cell types (observed probability mass). The proportion of mass derived from PBA and Stationary OT represent the expected proportion of cells ending up with a particular lineage fate, though individual cells may have the potential to acquire multiple fates. Agreement between observed and expected probabilities were used as a metric for evaluating performance of the probabilistic algorithms. Both PBA and Stationary OT inferred fate probabilities were within 5% of the observed proportion of mass for each lineage at each timepoint.
